# Proxy gene-by-environment Mendelian randomization study confirms a causal effect of maternal smoking on offspring birthweight, but little evidence of long-term influences on offspring health

**DOI:** 10.1101/601443

**Authors:** Qian Yang, Louise A C Millard, George Davey Smith

**Affiliations:** Medical Research Council Integrative Epidemiology Unit at the University of Bristol, Bristol, BS8 2BN, UK; Population Health Sciences, Bristol Medical School, University of Bristol, Bristol, BS8 2BN, UK; Intelligent Systems Laboratory, Department of Computer Science, University of Bristol, Bristol, BS8 1UB, UK

**Keywords:** gene × environment, Mendelian randomization, proxy, maternal smoking, pregnancy

## Abstract

**Objective:** To validate a novel proxy gene-by-environment (G×E) Mendelian randomization (MR) approach by replicating the previously established effect of maternal smoking heaviness in pregnancy on offspring birthweight, and then use GxE MR to investigate the effect of smoking heaviness in pregnancy on offspring health outcomes in later life and grandchild’s birthweight.

**Design:** A proxy G×E MR using participants’ genotype (i.e. rs16969968 in *CHRNA5*) as a proxy for their mother’s genotype.

**Setting:** UK Biobank.

**Participants:** 289,684 white British men and women aged 40-69 in UK Biobank.

**Main outcome measures:** Participants’ birthweight and later life outcomes (height, body mass index, lung function, asthma, blood pressure, age at menarche, years of education, fluid intelligence score, depression/anxiety, happiness), and birthweight of female participants’ first child.

**Results:** In our proof of principle analysis, each additional smoking-increasing allele was associated with a 0.018 (95% confidence interval (CI): −0.026, −0.009) kg lower birthweight in the “maternal smoking during pregnancy” stratum, but no meaningful effect (−0.002kg; 95% CI: −0.008, 0.003) in the “maternal non-smoking during pregnancy” stratum (interaction P-value=0.004). We found little evidence of an effect of maternal smoking heaviness on participants’ later life outcomes. We found the differences in associations of rs16969968 with grandchild’s birthweight between grandmothers who did versus did not smoke were heterogeneous (interaction P-value=0.042) among female participants who did (−0.020kg per allele; 95% CI: −0.044, 0.003) versus did not (0.007kg per allele; 95% CI: −0.005, 0.020) smoke in pregnancy.

**Conclusions:** Our study demonstrated how offspring genotype can be used to proxy for mothers’ genotype in G×E MR. We confirmed the previously established causal effect of maternal smoking on offspring birthweight but found little evidence of an effect on long-term health outcomes in the offspring. For grandchild’s birthweight, the effect of grandmother’s smoking heaviness in pregnancy may be modulated by maternal smoking status in pregnancy.

**WHAT IS ALREADY KNOWN TO THIS TOPIC:** Heavier maternal smoking in pregnancy causes lower offspring birthweight Maternal smoking in pregnancy is also associated with offspring outcomes in later life and grandchild’s birthweight, but it is not known whether these associations are causal Understanding the transgenerational causal effects of maternal smoking heaviness in pregnancy is important to inform public health policies

**WHAT THIS STUDY ADDS:** The proxy gene-by-environment Mendelian randomization approach can be used to explore maternal effects on offspring phenotypes when maternal genetic information is unavailable The approach confirmed the causal effect of smoking on offspring birthweight.

Maternal smoking status in pregnancy modulates the effect of grandmother’s smoking heaviness in pregnancy on grandchild’s birthweight, highlighting the importance of smoking cessation before pregnancy in each generation

## Introduction

The developmental origins of health and disease hypothesis proposes that early life experiences, including those in utero, can have long-term health effects, and maternal pregnancy exposures are important to long-term health of offspring (1). Heavier maternal smoking in pregnancy is known to be causally associated with lower offspring birthweight (2–6), but its other effects in offspring are less clear. Multivariable regression in observational data showed that heavier maternal smoking during pregnancy was associated with offspring being shorter (7) and more overweight/obese (8, 9), and having higher blood pressure (10), but had mixed associations with age at menarche (11), respiratory (12), cognitive (13), and mental health (14). Heavier maternal smoking in pregnancy has also been associated with higher grandchild’s birthweight in certain subpopulations (15–17). It is unclear whether these associations reflect a causal effect of maternal smoking in pregnancy, as they may be due to residual confounding. Some studies have assessed this using paternal smoking as a ‘negative control’ since an effect via uterine environment would be observed in mothers but not fathers, such that similar-magnitude associations would indicate confounding via shared familial, social, environmental and genetic factors (2, 5, 18). Negative control studies suggest little evidence of a causal effect on offspring body mass index (BMI) (2, 5, 8), blood pressure (19, 20) and depression (21).

Mendelian randomization (MR) provides an alternative way to explore this question by using single nucleotide polymorphisms (SNPs) as instrumental variables (IVs) for an exposure of interest. MR is less prone to confounding as germline genetic variants are randomly allocated at meiosis and not influenced by subsequent socioeconomic and health behaviours (22, 23). MR has been applied in a gene-by-environment (G×E) framework (24, 25), which requires variation in the strength of the gene-exposure association across strata of another factor. If there is a causal effect of the IV on the outcome via the exposure of interest, then we would expect the association of the IV with the outcome to vary in proportion to the gene-exposure association. The rs1051730/rs16969968 (*CHRNA5*) SNPs, previously robustly associated with smoking heaviness amongst smokers (26), have been widely used as IVs for smoking heaviness in GxE MR studies (3, 27–29). A causal effect of the smoking heaviness IV on an outcome should be seen amongst ever but not amongst never smokers if the effect is via smoking heaviness rather than other pathways (24, 25). G×E MR has also been used to assess cross-generational causal effects. A smoking heaviness IV has been associated with lower offspring birthweight amongst mothers who smoked in pregnancy but not amongst mothers who did not smoke in pregnancy, suggesting the genetic instrument affects birthweight through maternal smoking (3).

It is usually difficult to investigate transgenerational associations due to a lack of data across the generations of interest. Thus, previous work has sought to test transgenerational associations using available traits as proxies for unmeasured traits of interest. A Norwegian cohort aimed to examine whether women’s smoking in adulthood was related to their mothers’ smoking habits (that were not recorded) and hence used maternal smoking-related mortality as a proxy (30). Recently, a case-control by proxy approach has been proposed (31). Participants’ genotypes were used to proxy unavailable parental genotypes, and their associations were tested against parental diagnosis of Alzheimer’s disease in UK Biobank (31), since Alzheimer’s disease was much more prevalent in the parents than the participants (aged between 40 and 69 at baseline in 2006-2010 (32)). Our study aimed to demonstrate how an analogous approach can be used within a G×E MR framework to test maternal-offspring effects when maternal genotype is not available, using offspring genotype as a proxy for the maternal genotype. First, we performed a proof of principle analysis to demonstrate this approach, testing the previously established finding that maternal smoking in pregnancy leads to lower offspring birthweight. Second, we tested for causal effects of maternal smoking on offspring later life outcomes. Finally, we tested for a causal effect of grandmother’s smoking on grandchild’s birthweight.

## Methods

### Study population

Our study was conducted using UK Biobank, a population-based cohort of more than 500,000 men and women in the UK. This study collected a large and diverse range of data from physical measures, questionnaires and hospital episode statistics (32). Of 463,013 participants of European descent with genetic data passing initial quality control (i.e. genetic sex same as reported sex, XX or XY in sex chromosome and no outliers in heterozygosity and missing rates) (33), 289,684 participants (54% women) of white British descent were eligible for inclusion in our analyses (Supplementary Figure 1). We refer to the UK Biobank participants as generation one (G1), and their parents and offspring as G0 and G2, respectively.

### Genetic IV for maternal smoking

The rs16969968 SNP located in *CHRNA5* has been robustly associated with smoking heaviness (26). Ideally, we would use the maternal rs16969968 as an IV for the heaviness of maternal smoking, but in UK Biobank parental genetic data are not available. Hence, we used rs16969968 of the UK Biobank participants as a proxy for that of their mothers’, coded as the number of smoking heaviness increasing alleles.

### Smoking phenotypes

We used participants’ answers to the question “Did your mother smoke regularly around the time when you were born?” as a proxy for G0 smoking during pregnancy. Participants were also asked to report their smoking status (current/former/never). We derived a binary ever versus never measure of smoking status by combining current and former smokers. For female participants with at least one live birth, we derived a measure denoting whether they smoked during the pregnancy of their first child (see Supplementary Methods).

### Outcomes in participants (G1)

We used baseline data measured at the UK Biobank initial assessment center. Anthropometric traits included participants’ birthweight (kg, self-reported), standing height (cm) and BMI (kg/m^2^, constructed from standing height and weight). To assess lung function, forced vital capacity (L) and forced expiratory volume in 1-second (L) were measured by spirometry. Participants reported whether they had had asthma via the question “Has a doctor ever told you that you have had any of the following conditions?” (with an option of asthma) (34). Systolic and diastolic blood pressure (mmHg) were measured twice using a digital monitor or a manual sphygmomanometer if the digital monitor could not be employed, and we took the average of the two readings. Female participants reported their age at menarche. We derived years of education based on qualifications achieved by participants, as described previously (35). We included follow-up data of a subset of participants to define intelligence and depression/anxiety. Fluid intelligence score was generated as an unweighted sum of the number of correct answers given to 13 questions, and we used the earliest score if we had data at multiple time points (36). We defined depression/anxiety cases as participants that either answered “Yes” to “Have you ever seen a general practitioner (GP) for nerves, anxiety, tension or depression?” or “Have you ever seen a psychiatrist for nerves, anxiety, tension or depression?”, or had hospital episode coded using ICD-10 (37). Happiness was assessed via a question – “In general how happy are you?”, with six categories ranging from “extremely happy” to “extremely unhappy”.

### Outcomes in participants’ offspring (G2)

The female participants with at least one live birth were asked to report their first child’s birthweight. Male participants were not asked to report the birthweight of their offspring.

#### Statistical analyses

##### Proof of principle analysis: testing the causal effect of maternal (G0) smoking heaviness in pregnancy on participants’ (G1) birthweight

In this proof of principle analysis, we seek to replicate the finding, previously established using GxE MR and many other methods (6), that heavier maternal smoking causes lower offspring birthweight. We use our proxy GxE approach, where participants’ (G1) genotype is used as a proxy for their mothers’ (G0) genotype. To assess whether rs16969968 affects participants’ birthweight via G0 smoking in pregnancy, we stratified our G1 sample by G0 smoking status during pregnancy, and then tested the associations of rs16969968 with birthweight in each stratum using multivariable linear regression. Since birth precedes smoking initiation, participants’ genotype cannot affect birthweight through their own smoking heaviness, which means we do not need to consider smoking status of participants (Figure 1A). We included participants’ sex as a covariate to reduce variation in their birthweight and the first ten principal components to control for population stratification. We assumed an additive genetic effect and identified the strength of interaction between strata using Cochran’s Q test for heterogeneity.

**Figure 1.**
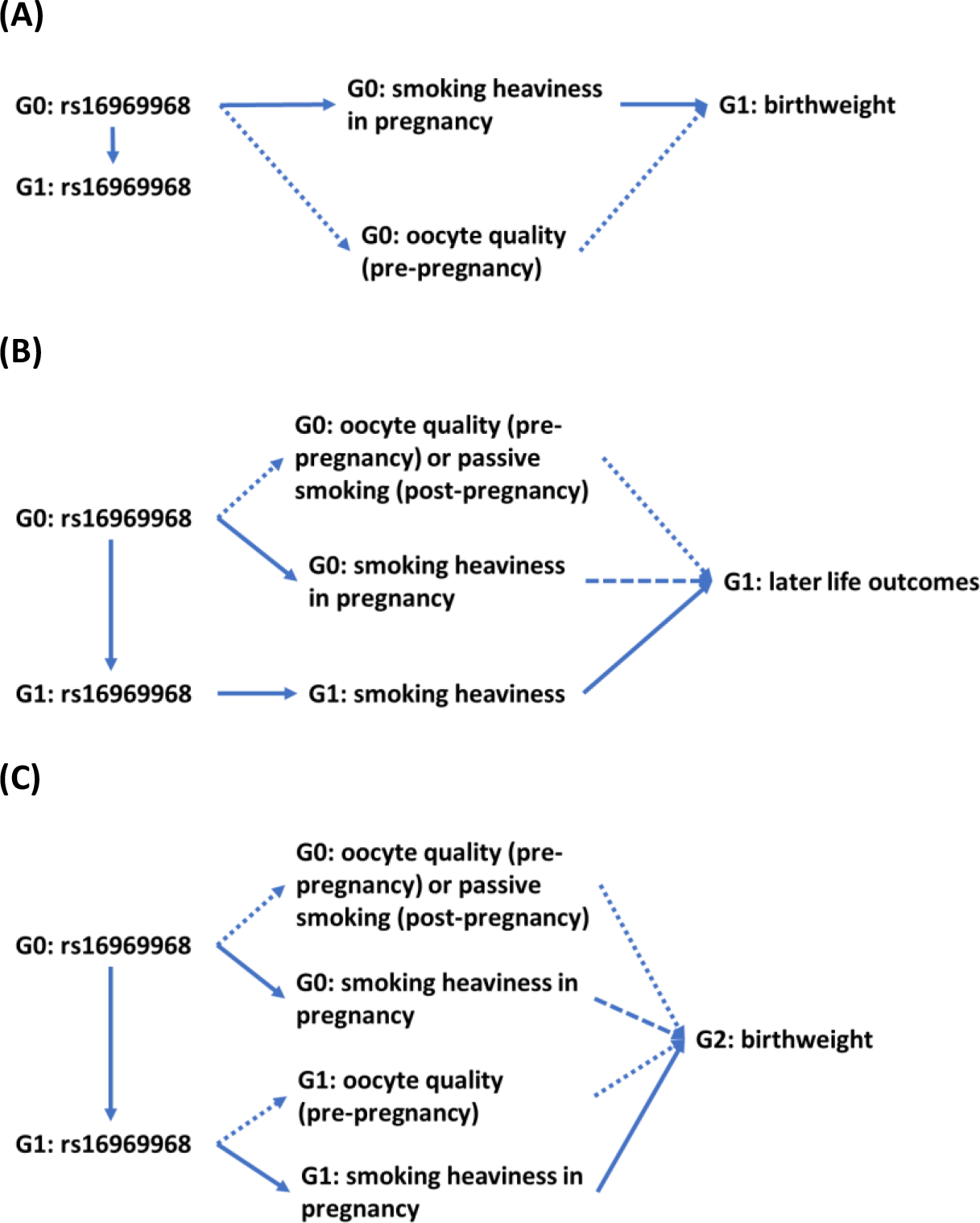
Directed acyclic graphs (DAGs) of this study. Generation (G)0: UK Biobank participants’ mother; G1: UK Biobank participants themselves; G2: First offspring of UK Biobank participants. A) Assessing the effect of G0 smoking heaviness on G1 birthweight: We used G1 rs16969968 as a proxy for G0 rs16969968 and stratified on G0 smoking status in pregnancy. There is no backdoor path (54) via G1 smoking heaviness since G1 cannot smoke before they were born. Maternal smoking outside of pregnancy might influence the outcome (52), e.g. via oocyte quality, causing an alternate path between rs16969968 and G1 birthweight (shown as 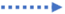). B) Assessing the effect of G0 smoking on G1 later life outcomes: Besides the paths described in (A), there is a backdoor path from G1 rs16969968 via G1 life-course smoking heaviness to the outcomes. To estimate the effect of G0 smoking heaviness in pregnancy (shown as 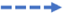), we need to block this backdoor path by further stratifying on G1 smoking status. C) Assessing the effect of G0 smoking on G2 birthweight: Besides the paths described in (A), there is a backdoor path from G1 rs16969968 via G1 smoking heaviness in pregnancy to the outcomes. To estimate the effect of G0 smoking heaviness in pregnancy (shown as 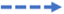), we need to block this backdoor path by further stratifying on G1 smoking status in pregnancy. G1 pre-pregnancy smoking might influence G2 birthweight (shown as 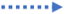). See further DAGs in the Supplementary Figure 2 illustrating potential sources of bias due to conditioning on a collider.

##### Testing for causal effects of G0 smoking in pregnancy on G1 later life outcomes

We use the proxy GxE MR approach to test for causal effects of maternal (G0) smoking heaviness on offspring (G1) height, BMI, lung function, asthma, blood pressure, age at menarche, education, intelligence, depression/anxiety and happiness. In contrast to our proof of principle example where participants smoking in adulthood cannot influence their birthweight, participants’ rs16969968 could affect these outcomes via both maternal (G0) and participants’ (G1) smoking heaviness (Figure 1B). To assess whether rs16969968 may affect these outcomes via maternal versus participants’ smoking, we stratified on both maternal and participants’ smoking status. In each stratum, we examined associations of rs16969968 with height, BMI, lung function, blood pressure, age at menarche, education and intelligence using linear regression, asthma and depression/anxiety using logistic regression, and happiness using ordinal logistic regression. We included participants’ age at baseline, sex and the first ten genetic principal components as covariates.

Height and age at menarche manifest around the time of puberty such that participants’ own smoking can only affect these if they started smoking before these outcomes are determined. We conducted sensitivity analyses for these outcomes stratifying G1 participants according to whether they were ever smokers before achieving their adulthood height (assuming age at 17 for men and 15 for women (38)) or their age at menarche.

##### Testing for causal effects of G0 smoking in pregnancy on grandchild’s (G2) birthweight

To test for a causal effect of participants mothers’ smoking on birthweight of participants’ offspring, we stratified G1 women based on their own and their mothers’ smoking status during pregnancy, as rs16969968 could affect G2 birthweight through both G0 and G1 smoking heaviness (Figure 1C). Within each stratum, we assessed associations of rs16969968 with G2 birthweight using linear regression, adjusting for the first ten genetic principal components. We estimated the strength of interaction between G0 smokers and G0 non-smokers within each G1 stratum. We also calculated a difference (39) in those associations between G0 smokers and G0 non-smokers within each G1 stratum, and estimated the strength of interaction between two differences to investigate whether G1 smoking status modulates the effect of rs16969968 on G2 birthweight.

Our G × E MR may be vulnerable to collider bias (29, 40, 41), as we stratified on smoking status (Supplementary Figure 2). Therefore, we tested associations of rs16969968 with G0 and G1 smoking status and potential confounders available in UK Biobank (see Supplementary Methods). We also tested observational associations of maternal (G0) smoking status with offspring (G1) smoking status and all outcomes for comparison with our MR results. Analyses were performed using R version 3.5.1 (R Foundation for Statistical Computing, Vienna, Austria).

##### Patient and public involvement

The current research was not informed by patient and public involvement because it used secondary data. However, future research following on from our findings should be guided by patient and public opinions.

## Results

Characteristics of participants across sex are shown in Supplementary Table 1. Each additional smoking-increasing allele of participants’ rs16969968 was associated with a 1.02 (95% confidence interval [CI]: 1.01, 1.03; P-value = 5×10^−3^) higher odds of their mothers’ smoking in pregnancy, a 0.98 (95% CI: 0.97, 0.99; P-value = 7×10^−4^) lower odds of being an ever (versus never) smoker themselves, and a 1.06 (95% CI: 1.04, 1.09; P-value = 3×10^−7^) higher odds that female participants were a smoker (versus non-smoker) in their own pregnancy. We found little evidence of an association between rs16969968 and potential confounders, with small associations for participants’ age and years of education in some strata (Supplementary Table 2).

Our proof of principle analysis found that, amongst participants whose mothers smoked in pregnancy, each additional smoking-increasing allele was associated with a 0.018kg lower birthweight (95% CI: −0.026, −0.009) after adjustment for covariates (Figure 2). Amongst participants whose mothers did not smoke in pregnancy, we found little evidence for an association of rs16969968 with birthweight (−0.002kg [95% CI: −0.008, 0.003]), and we observed heterogeneity between these associations (interaction P-value = 0.004).

**Figure 2.**
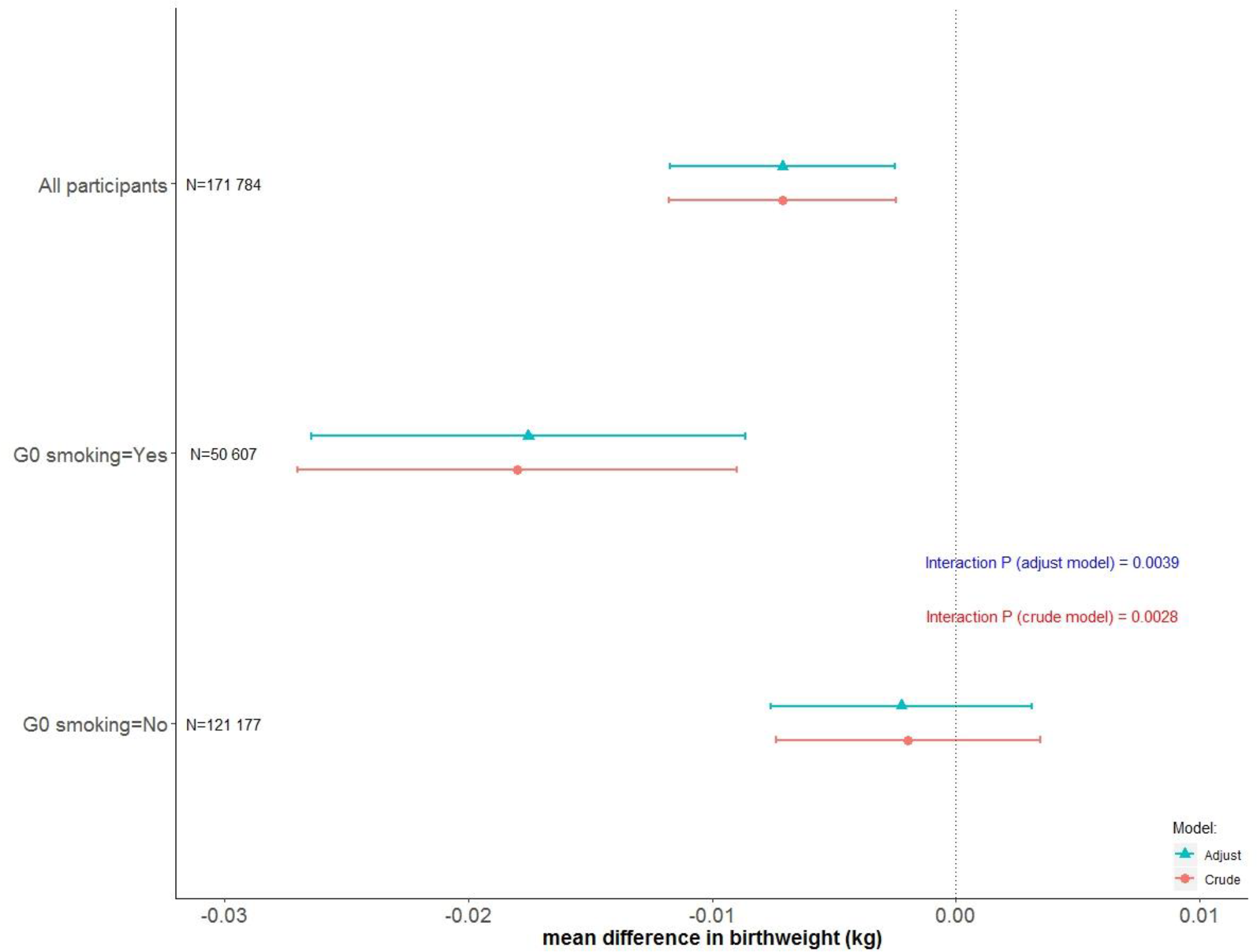
The associations of rs16969968 of UK Biobank participants with their own birthweight by their mothers’ smoking status during pregnancy. Generation (G)0: UK Biobank participants’ mother; G1: UK Biobank participants themselves. Estimates are the mean difference of G1 birthweight per each smoking-heaviness increasing allele of rs16969968. Associations are adjusted for sex of participants and the first ten principal components. The number of participants was listed for each analysis.

Figure 3 showed estimates of rs16969968 on the 12 outcomes in the UK Biobank participants. Overall, within each stratum, the estimates were broadly consistent between those whose mothers smoked and those whose mothers did not, except for height among participants who never smoked (all interaction P-values were in Supplementary Table 3). Each additional smoking-increasing allele was associated with a 0.115cm lower height (95% CI: −0.200, −0.030) among never smokers whose mothers smoked in pregnancy, but a 0.002cm lower height (95% CI: −0.057, 0.053) among never smokers whose mothers did not smoke in pregnancy (interaction P-value = 0.029). However, this difference was not observed amongst ever smokers (Figure 3A). We obtained largely consistent results in sensitivity analyses (Supplementary Figure 3).

**Figure 3.**
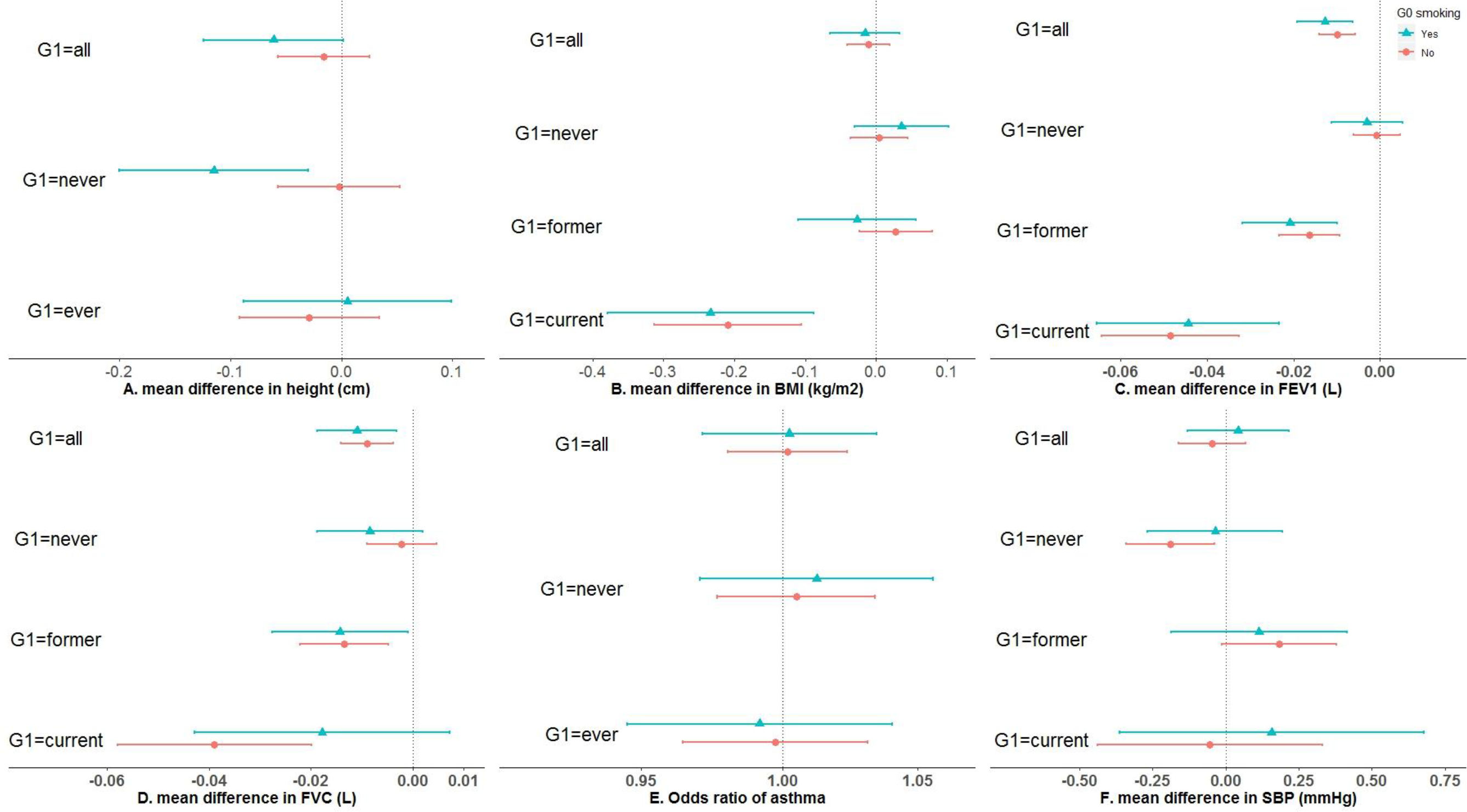
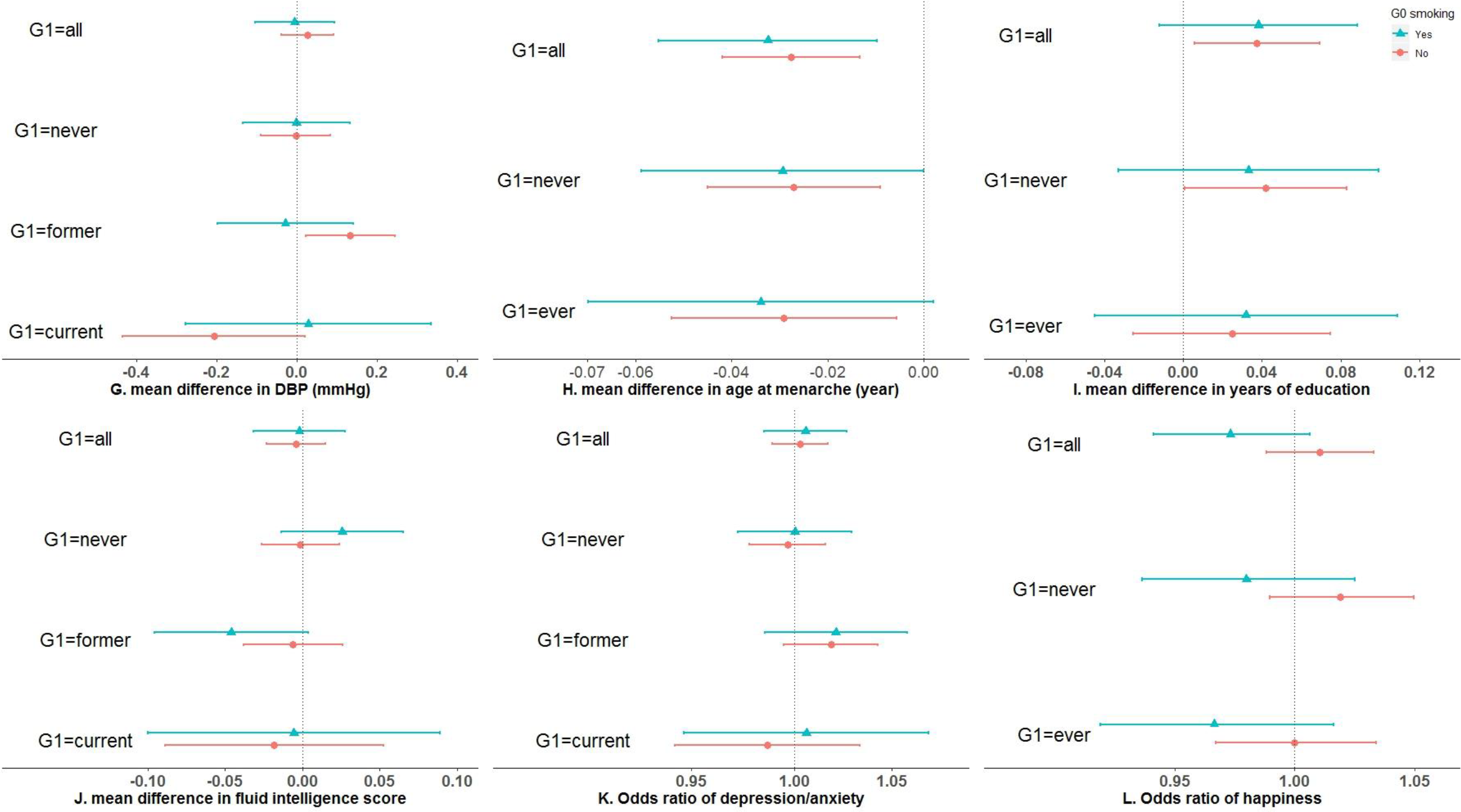
The associations of rs16969968 with 12 outcomes in UK Biobank participants by their mothers’ smoking status during pregnancy and their own smoking status. Generation (G)0: UK Biobank participants’ mother; G1: UK Biobank participants themselves. Estimates are the mean difference (or change in odds) of G1 outcome per each smoking-heaviness increasing allele of rs16969968. We adjusted for age and sex of participants for outcomes except for menarche, and the first ten principal components for all 12 outcomes. We combined G1 current and former smokers into ever smokers for height, menarche, education, asthma and happiness to enlarge sample sizes given smoking cessation may not have a rapid impact on them. Abbreviations: BMI, body mass index, DBP, diastolic blood pressure; FEV_1_, forced expiratory volume in 1-second; FVC, forced vital capacity; SBP, systolic blood pressure.

Figure 4 showed estimates of rs16969968 on grandchild’s birthweight. Among female participants who did not smoke in pregnancy, each additional smoking-increasing allele was associated with a 0.007kg higher grandchild’s birthweight difference (95% CI: −0.005, 0.020) between grandmothers who did versus did not smoke in pregnancy. However, this difference was −0.020kg per allele (95% CI: −0.044, 0.003) among female participants who smoked in pregnancy. These two differences were heterogeneous (−0.028kg per allele [95% CI: −0.055, −0.001]; interaction P-value=0.042).

**Figure 4.**
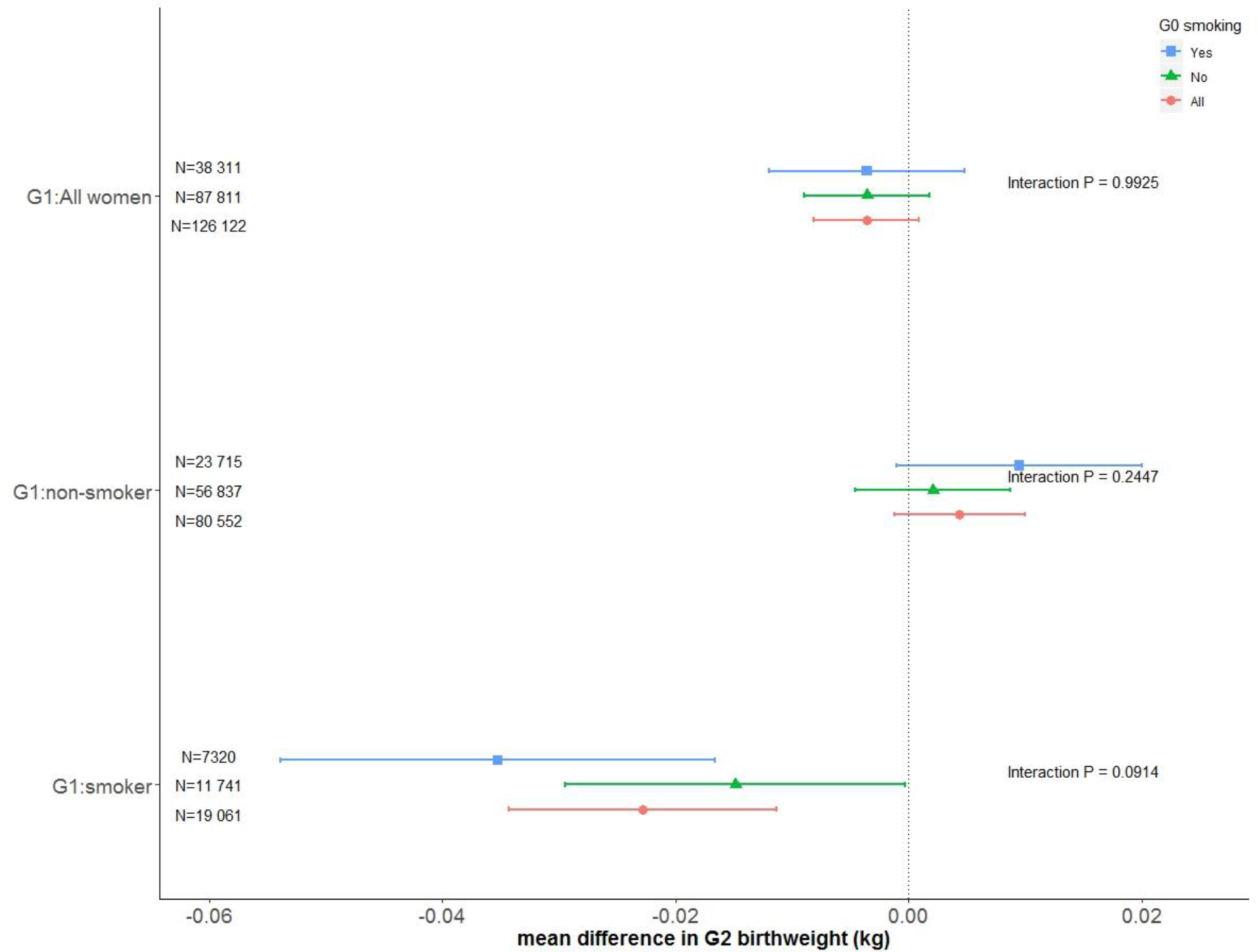
The associations of rs16969968 of UK Biobank women participants with their first child’s birthweight by their mothers’ and their own smoking status during pregnancy, after adjusting for the first ten genetic principal components. Generation (G)0: UK Biobank participants’ mother; G1: UK Biobank participants themselves; G2: First offspring of UK Biobank participants. Estimates are the mean difference of G2 birthweight per each smoking-heaviness increasing allele of rs16969968. Interactions are tested between G0 smokers (blue line) and non-smokers (green line) with their P-values presented. All women in G1 included G1 smokers, G1 non-smokers and G1 women whose smoking status in pregnancy was missing.

The directions of observational estimates were consistent with our MR estimates for both participants’ and their child’s birthweight. Our observational analyses also found associations of maternal smoking in pregnancy with offspring later life outcomes, where smoking in pregnancy was associated with lower height, higher BMI, poorer lung function, higher risk of asthma, earlier age at menarche, higher blood pressure, and poorer cognitive and mental health (Supplementary Table 4).

## Discussion

### Principle findings and comparison with the literature

In this study, we have demonstrated how G×E MR can be used to test transgenerational causal effects of maternal smoking heaviness in pregnancy using participants’ genotype as a proxy for their mothers’ genotype. Our proof of principle analysis identified an effect of heavier maternal smoking on lower offspring birthweight, consistent with previous studies (2–6). Our MR study also confirmed previously established causal effects of participants’ smoking on their own health, where heavier smoking reduced BMI (27) and lead to impaired lung function (42), but found little evidence of an effect on asthma risk (43) or blood pressure (28).

Our tests of effects of maternal smoking heaviness on offspring later life health outcomes were not conclusive, given a lack of precision for many of our MR estimates. We found little evidence of an effect on BMI, lung function, asthma, blood pressure, cognition, depression/anxiety or happiness. These findings were consistent with negative control studies for BMI (2, 8), blood pressure (19, 20) and depression/anxiety (21), although our estimation of interactions is not directly quantitatively comparable to their estimation of effects of ever/never smoking or smoking heaviness categories in observational studies. Our MR results found little evidence to support findings from our own and previous observational studies indicating maternal smoking led to poorer lung function (44), higher risk of asthma (45, 46), and lower happiness in offspring (47). This may be due to residual confounding in observational associations, or because of low statistical power in MR. Previous studies did not use the same cognition measurement approaches as used UK Biobank, making our results for this outcome less comparable. We observed lower offspring adulthood height according to maternal smoking in never smokers but not in ever smokers, which could be a chance finding given we tested multiple outcomes.

We found little evidence of an effect of maternal smoking in pregnancy on offspring age at menarche. However, we did find an effect of rs16969968 on age at menarche across strata of smoking status (of both the participant and participants’ mother) suggesting that rs16969968 may have horizontal pleiotropic effects on age at menarche (e.g. via smoking outside of pregnancy). Future MR studies could examine this (25).

Our observational results were consistent with previous observational studies (15–17) by showing a positive association of grandmother’s (G0) smoking in pregnancy with grandchild’s (G2) birthweight after adjusting for mother’s (G1) smoking in pregnancy. Although our G×E MR was vulnerable to insufficient statistical power, we did find evidence that female G1 smoking in pregnancy modulates the effect of G0 smoking heaviness in pregnancy on G2 birthweight, consistent with previous observational findings (15–17). These results highlight the importance of both grandmother’s and maternal smoking in pregnancy for fetal growth, which could have implications for public health interventions aiming to reduce the prevalence of low birthweight.

### Strength of weakness of this study

We now discuss some limitations of this work. First, our proxy G×E MR used offspring genotype as a proxy for maternal genotype and offspring rs16969968 contains 50% information from fathers. This may cause regression dilution bias in each stratum, where the measurement error in the SNP biases associations towards the null (48). However, we checked the extent that this might affect our results, by comparing the associations of participant’s rs16969968 with their own birthweight versus their child’s birthweight for smokers during pregnancy, and found little difference (−0.005kg (95% CI: −0.020, 0.009)) between them. Second, we stratified on smoking status which rs16969968 was weakly associated with. Stratification on colliders (between rs16969968 and outcomes) may bias our MR estimates (see Supplementary Figure 2) (40, 41). Additionally, we used a highly selected sample related to smoking (49) and had missing data in outcomes. These may also make our MR estimates vulnerable to selection bias (50). However, previous evidence (29, 51) and our genetic associations with measured confounders indicated that these selection effects may not be large enough to have a considerable impact on our MR estimates. Third, rs16969968 predicts life-course smoking heaviness and not just in pregnancy. Women who smoked in pregnancy may also smoke outside of pregnancy. Therefore, the effect of maternal smoking might be via other pathways such as poor oocyte quality for offspring birthweight, or postnatal maternal smoking (e.g. passive smoke exposure) for adulthood outcomes among offspring (52).

Fourth, both participants’ and their mothers’ (G0) smoking status may be misclassified. Participants were asked to report whether their mother smoked around the time of their birth and we used this as our measure of G0 smoking in pregnancy. This means that G0 smokers might have smoked during all their pregnancy, part of their pregnancy or started smoking shortly after giving birth. Effects of smoking heaviness in pregnancy may vary according to the duration and pregnancy period during which a woman smoked. For instance, previous work found that smoking in the first trimester was not associated with lower birthweight in offspring suggesting that later stages may be more important for fetal growth (3, 15). Similarly, participants reported their smoking status at baseline, but this may not reflect their smoking status at an important time point for a given outcome. For, instance, participants’ height and age at menarche can only be affected by their own smoking behaviour if they started smoking before achieving adult height or the onset of puberty. We performed sensitivity analyses for height and age at menarche using estimates of participants smoking status before these outcomes. For height, this assumed that men and women achieved their adult height at 17 and 15 years old (38), respectively, as this information was not available in UK Biobank. Fifth, we tested several hypotheses which increases the probability that our identified associations may be due to chance. Finally, our study may lack statistical power due to small sample sizes in strata and the low power of tests for interactions (53). We were unable to account for grandchild’s sex in our models assessing the impact of grandmother’s smoking in pregnancy since that is unavailable in UK Biobank, which may also reduce our statistical power. MR studies with larger sample sizes and hence greater statistical power are needed to further investigate transgenerational effects of smoking heaviness, together with studies in which both maternal and offspring genotype are known.

### Conclusion

G×E MR demonstrates how offspring genotype can be used to proxy for maternal genotype to investigate causal effects of maternal smoking heaviness in pregnancy when maternal genotype is unavailable. We demonstrated our proxy GxE approach by replicating the previously identified effect of heavier smoking on lower offspring birthweight. We found little evidence of a causal effect of maternal smoking heaviness on offspring’s later life outcomes. Finally, we found evidence that the effect of grandmother’s smoking in pregnancy on grandchild’s birthweight may be modulated by mother’s smoking status in pregnancy. Further studies with larger sample sizes are needed to improve statistical power.

## Acknowledgments

This research has been conducted using the UK Biobank Resource under Application Number 16729 (dataset ID 11148 and 21753).

## Footnotes

### Contributors

QY contributed to the design of the study, performed all analyses, wrote the first version of the manuscript, critically reviewed and revised the manuscript and approved the final version of the manuscript as submitted. LACM contributed to the design of the study, critically reviewed and revised the manuscript and approved the final version of the manuscript as submitted. GDS conceptualized the study, contributed to the design of the study, critically reviewed and revised the manuscript and approved the final version of the manuscript as submitted. GDS is the guarantor. The corresponding author attests that all listed authors meet authorship criteria and that no others meeting the criteria have been omitted.

### Ethical approval

The UK Biobank received ethical approval from the research ethics committee (REC reference for UK Biobank 11/NW/0382) and participants provided written informed consent.

### Data sharing

The data reported in this paper are available by applying directly to the UK Biobank. All code used to produce the results can be accessed at https://github.com/MRCIEU/MR-maternal-smoking. Git tag v0.1 corresponds to the version presented here.

### Transparency

The lead author (the manuscript’s guarantor) affirms that this manuscript is an honest, accurate, and transparent account of the study being reported; that no important aspects of the study have been omitted; and that any discrepancies from the study as originally planned (and, if relevant, registered) have been explained.

This is an Open Access article distributed in accordance with the terms of the Creative Commons Attribution (CC BY 4.0) license, which permits others to distribute, remix, adapt and build upon this work, for commercial use, provided the original work is properly cited.

See: http://creativecommons.org/licenses/by/4.0/.

## References

1. Sharp GC, Lawlor DA, Richardson SS. It’s the mother!: How assumptions about the causal primacy of maternal effects influence research on the developmental origins of health and disease. Soc Sci Med 2018;213:20–7. doi: 10.1016/j.socscimed.2018.07.035

2. Davey Smith G. Negative control exposures in epidemiologic studies. Epidemiology 2012;23:350–1; author reply 1-2. doi: 10.1097/EDE.0b013e318245912c

3. Tyrrell J, Huikari V, Christie JT, et al. Genetic variation in the 15q25 nicotinic acetylcholine receptor gene cluster (CHRNA5-CHRNA3-CHRNB4) interacts with maternal self-reported smoking status during pregnancy to influence birth weight. Hum Mol Genet 2012;21:5344–58. doi: 10.1093/hmg/dds372

4. Rice F, Harold GT, Boivin J, Hay DF, van den Bree M, Thapar A. Disentangling prenatal and inherited influences in humans with an experimental design. Proc Natl Acad Sci U S A 2009;106:2464–7. doi: 10.1073/pnas.0808798106

5. Davey Smith G. Assessing intrauterine influences on offspring health outcomes: can epidemiological studies yield robust findings? Basic Clin Pharmacol Toxicol 2008;102:245–56. doi: 10.1111/j.1742-7843.2007.00191.x

6. Krieger N, Davey Smith G. The tale wagged by the DAG: broadening the scope of causal inference and explanation for epidemiology. Int J Epidemiol 2016;45:1787–808. doi: 10.1093/ije/dyw114

7. Howe LD, Matijasevich A, Tilling K, et al. Maternal smoking during pregnancy and offspring trajectories of height and adiposity: comparing maternal and paternal associations. Int J Epidemiol 2012;41:722–32. doi: 10.1093/ije/dys025

8. Riedel C, Schonberger K, Yang S, et al. Parental smoking and childhood obesity: higher effect estimates for maternal smoking in pregnancy compared with paternal smoking--a meta-analysis. Int J Epidemiol 2014;43:1593–606. doi: 10.1093/ije/dyu150

9. Albers L, Sobotzki C, Kuss O, et al. Maternal smoking during pregnancy and offspring overweight: is there a dose-response relationship? An individual patient data meta-analysis. Int J Obes (Lond) 2018;42:1249–64. doi: 10.1038/s41366-018-0050-0

10. Brion MJ, Leary SD, Lawlor DA, Davey Smith G, Ness AR. Modifiable maternal exposures and offspring blood pressure: a review of epidemiological studies of maternal age, diet, and smoking. Pediatr Res 2008;63:593–8. doi: 10.1203/PDR.0b013e31816fdbd3

11. Chen Y, Liu Q, Li W, Deng X, Yang B, Huang X. Association of prenatal and childhood environment smoking exposure with puberty timing: a systematic review and meta-analysis. Environ Health Prev Med 2018;23:33. doi: 10.1186/s12199-018-0722-3

12. Jayes L, Haslam PL, Gratziou CG, et al. SmokeHaz: Systematic Reviews and Meta-analyses of the Effects of Smoking on Respiratory Health. Chest 2016;150:164–79. doi: 10.1016/j.chest.2016.03.060

13. Clifford A, Lang L, Chen R. Effects of maternal cigarette smoking during pregnancy on cognitive parameters of children and young adults: a literature review. Neurotoxicol Teratol 2012;34:560–70. doi: 10.1016/j.ntt.2012.09.004

14. Tiesler CM, Heinrich J. Prenatal nicotine exposure and child behavioural problems. Eur Child Adolesc Psychiatry 2014;23:913–29. doi: 10.1007/s00787-014-0615-y

15. Ding M, Yuan C, Gaskins AJ, et al. Smoking during pregnancy in relation to grandchild birth weight and BMI trajectories. PLoS One 2017;12:e0179368. doi: 10.1371/journal.pone.0179368

16. Miller LL, Pembrey M, Davey Smith G, Northstone K, Golding J. Is the growth of the fetus of a non-smoking mother influenced by the smoking of either grandmother while pregnant? PLoS One 2014;9:e86781. doi: 10.1371/journal.pone.0086781

17. Hypponen E, Davey Smith G, Power C. Effects of grandmothers’ smoking in pregnancy on birth weight: intergenerational cohort study. BMJ 2003;327:898. doi: 10.1136/bmj.327.7420.898

18. Macdonald-Wallis C, Tobias JH, Davey Smith G, Lawlor DA. Parental smoking during pregnancy and offspring bone mass at age 10 years: findings from a prospective birth cohort. Osteoporos Int 2011;22:1809–19. doi: 10.1007/s00198-010-1415-y

19. Brion MJ, Leary SD, Davey Smith G, Ness AR. Similar associations of parental prenatal smoking suggest child blood pressure is not influenced by intrauterine effects. Hypertension 2007;49:1422–8. doi: 10.1161/hypertensionaha.106.085316

20. Leary SD, Brion MJ, Lawlor DA, Smith GD, Ness AR. Lack of emergence of associations between selected maternal exposures and offspring blood pressure at age 15 years. J Epidemiol Community Health 2013;67:320–6. doi: 10.1136/jech-2012-201784

21. Taylor AE, Carslake D, de Mola CL, et al. Maternal Smoking in Pregnancy and Offspring Depression: a cross cohort and negative control study. Sci Rep 2017;7:12579. doi: 10.1038/s41598-017-11836-3

22. Davey Smith G, Lawlor DA, Harbord R, Timpson N, Day I, Ebrahim S. Clustered environments and randomized genes: a fundamental distinction between conventional and genetic epidemiology. PLoS Med 2007;4:e352. doi: 10.1371/journal.pmed.0040352

23. Davey Smith G, Ebrahim S. ‘Mendelian randomization’: can genetic epidemiology contribute to understanding environmental determinants of disease? Int J Epidemiol 2003;32:1–22. doi:

24. Davey Smith G. Use of genetic markers and gene-diet interactions for interrogating population-level causal influences of diet on health. Genes Nutr 2011;6:27–43. doi: 10.1007/s12263-010-0181-y

25. Spiller W, Slichter D, Bowden J, Davey Smith G. Detecting and correcting for bias in Mendelian randomization analyses using Gene-by-Environment interactions. Int J Epidemiol 2018. doi: 10.1093/ije/dyy204

26. Munafo MR, Timofeeva MN, Morris RW, et al. Association between genetic variants on chromosome 15q25 locus and objective measures of tobacco exposure. J Natl Cancer Inst 2012;104:740–8. doi: 10.1093/jnci/djs191

27. Freathy RM, Kazeem GR, Morris RW, et al. Genetic variation at CHRNA5-CHRNA3-CHRNB4 interacts with smoking status to influence body mass index. Int J Epidemiol 2011;40:1617–28. doi: 10.1093/ije/dyr077

28. Linneberg A, Jacobsen RK, Skaaby T, et al. Effect of Smoking on Blood Pressure and Resting Heart Rate: A Mendelian Randomization Meta-Analysis in the CARTA Consortium. Circ Cardiovasc Genet 2015;8:832–41. doi: 10.1161/circgenetics.115.001225

29. Millard LAC, Munafo M, Tilling K, Wootton RE, Davey Smith G. MR-pheWAS with stratification and interaction: Searching for the causal effects of smoking heaviness identified an effect on facial aging. bioRxiv 2018. doi: 10.1101/441907

30. Kvalvik LG, Skjaerven R, Klungsoyr K, Vollset SE, Haug K. Can ‘early programming’ be partly explained by smoking? Results from a prospective, population-based cohort study. Paediatr Perinat Epidemiol 2015;29:50–9. doi: 10.1111/ppe.12164

31. Liu JZ, Erlich Y, Pickrell JK. Case-control association mapping by proxy using family history of disease. Nat Genet 2017;49:325–31. doi: 10.1038/ng.3766

32. Sudlow C, Gallacher J, Allen N, et al. UK biobank: an open access resource for identifying the causes of a wide range of complex diseases of middle and old age. PLoS Med 2015;12:e1001779. doi: 10.1371/journal.pmed.1001779

33. Mitchell R, Hemani G, Dudding T, Paternoster L. UK Biobank Genetic Data: MRC-IEU Quality Control, Version 1. 06 Nov 2017. https://data.bris.ac.uk/data/dataset/3074krb6t2frj29yh2b03x3wxj.

34. Zhu Z, Lee PH, Chaffin MD, et al. A genome-wide cross-trait analysis from UK Biobank highlights the shared genetic architecture of asthma and allergic diseases. Nat Genet 2018;50:857–64. doi: 10.1038/s41588-018-0121-0

35. Okbay A, Beauchamp JP, Fontana MA, et al. Genome-wide association study identifies 74 loci associated with educational attainment. Nature 2016;533:539–42. doi: 10.1038/nature17671

36. Hill WD, Marioni RE, Maghzian O, et al. A combined analysis of genetically correlated traits identifies 187 loci and a role for neurogenesis and myelination in intelligence. Mol Psychiatry 2019;24:169–81. doi: 10.1038/s41380-017-0001-5

37. Howard DM, Adams MJ, Shirali M, et al. Genome-wide association study of depression phenotypes in UK Biobank identifies variants in excitatory synaptic pathways. Nat Commun 2018;9:1470. doi: 10.1038/s41467-018-03819-3

38. Tanner JM, Whitehouse RH, Takaishi M. Standards from birth to maturity for height, weight, height velocity, and weight velocity: British children, 1965. In Arch Dis Child 1966;41:454–71. doi:

39. Knezevic A. StatNews # 73: Overlapping Confidence Intervals and Statistical Significance. Oct 2008;. https://www.cscu.cornell.edu/news/statnews/stnews73.pdf.

40. Munafo MR, Tilling K, Taylor AE, Evans DM, Davey Smith G. Collider scope: when selection bias can substantially influence observed associations. Int J Epidemiol 2018;47:226–35. doi: 10.1093/ije/dyx206

41. Taylor AE, Davies NM, Ware JJ, VanderWeele T, Smith GD, Munafo MR. Mendelian randomization in health research: using appropriate genetic variants and avoiding biased estimates. Econ Hum Biol 2014;13:99–106. doi: 10.1016/j.ehb.2013.12.002

42. Colak Y, Afzal S, Lange P, Nordestgaard BG. Smoking, Systemic Inflammation, and Airflow Limitation: A Mendelian Randomization Analysis of 98 085 Individuals from the General Population. Nicotine Tob Res 2018. doi: 10.1093/ntr/nty077

43. Skaaby T, Taylor AE, Jacobsen RK, et al. Investigating the causal effect of smoking on hay fever and asthma: a Mendelian randomization meta-analysis in the CARTA consortium. Sci Rep 2017;7:2224. doi: 10.1038/s41598-017-01977-w

44. Magnus MC, Henderson J, Tilling K, Howe LD, Fraser A. Independent and combined associations of maternal and own smoking with adult lung function and COPD. Int J Epidemiol 2018;47:1855–64. doi: 10.1093/ije/dyy221

45. Silvestri M, Franchi S, Pistorio A, Petecchia L, Rusconi F. Smoke exposure, wheezing, and asthma development: a systematic review and meta-analysis in unselected birth cohorts. Pediatr Pulmonol 2015;50:353–62. doi: 10.1002/ppul.23037

46. Accordini S, Calciano L, Johannessen A, et al. A three-generation study on the association of tobacco smoking with asthma. Int J Epidemiol 2018;47:1106–17. doi: 10.1093/ije/dyy031

47. Menezes AM, Murray J, Laszlo M, et al. Happiness and depression in adolescence after maternal smoking during pregnancy: birth cohort study. PLoS One 2013;8:e80370. doi: 10.1371/journal.pone.0080370

48. Hutcheon JA, Chiolero A, Hanley JA. Random measurement error and regression dilution bias. Bmj 2010;340:c2289. doi: 10.1136/bmj.c2289

49. Swanson JM. The UK Biobank and selection bias. Lancet 2012;380:110. doi: 10.1016/s0140-6736(12)61179-9

50. Hughes RA, Davies NM, Davey Smith G, Tilling K. Selection bias in instrumental variable analyses. BioRxiv 2018. doi: 10.1101/192237

51. Gkatzionis A, Burgess S. Contextualizing selection bias in Mendelian randomization: how bad is it likely to be? Int J Epidemiol 2018. doi: 10.1093/ije/dyy202

52. Lawlor D, Richmond R, Warrington N, et al. Using Mendelian randomization to determine causal effects of maternal pregnancy (intrauterine) exposures on offspring outcomes: Sources of bias and methods for assessing them. Wellcome Open Res 2017;2:11. doi: 10.12688/wellcomeopenres.10567.1

53. Marshall SW. Power for tests of interaction: effect of raising the Type I error rate. Epidemiol Perspect Innov 2007;4:4. doi: 10.1186/1742-5573-4-4

54. Pearl J, Glymour M, Jewell NP. Causal inference in statistics A Primer: John Wiley & Sons Ltd, 2016.

