## Supplementary for "Proxy gene-by-environment Mendelian randomization study confirms a causal effect of maternal smoking on offspring birthweight, but little evidence of long-term influences on offspring health"

### SUPPLEMENTARY MATERIAL

Supplementary Methods

Supplementary Figure 1. Participant flow diagram

Supplementary Figure 2. Directed acyclic graphs illustrating potential collider bias

Supplementary Figure 3. The associations of rs16969968 with UK Biobank participants height and age at menarche in sensitivity analyses

Supplementary Figure 4. Illustration of edge cases when deriving participants smoking in pregnancy

Supplementary Table 1. Characteristics of the UK Biobank participants by sex

Supplementary Table 2. Associations of rs16969968 with potential confounders by strata of smoking status in participants (G1) and their mothers (G0), adjusted for the first ten principal components

Supplementary Table 3. Differences in associations of rs16969968 with 12 outcomes in participants (G1) across maternal (G0) smoking status in pregnancy

Supplementary Table 4. Observational associations of participants mothers' smoking in pregnancy with participants' smoking status and all outcomes

Supplementary Table 5. UK Biobank data field of variables used in this study

### SUPPLEMENTARY METHODS

*Smoking phenotypes*

We derived a measure denoting whether female participants who had at least one live birth smoked during the pregnancy of their first child, using their self-reported age at their first live birth and ages they reported starting and stopping smoking. As these details were recorded as a whole number of years, it was not always possible to determine whether a woman was a smoker during the pregnancy (illustrated in Supplementary Figure 4). For example, a participant reporting her first live birth occurring at 25 years old could have been pregnant at 25 years old only, or both 24 and 25 years old. If she was a current smoker and started smoking at 25 years old, it was unknown if this was after or during her pregnancy (Supplementary Figure 4A). If she was a current smoker and started smoking at 24 years old, she smoked at least in her late pregnancy. Similarly, if she stopped smoking at 24 or 25 years old (Supplementary Figure 4B&C), it was unknown if this was before or during their pregnancy. If she stopped smoking at 23 years old, she did not smoke during her pregnancy. Women who gave birth to their first child at age *b* were assigned as non-smokers in pregnancy if they: 1) never smoked, 2) currently smoked but started smoking when *b+1* years old or later, 3) formerly smoked but started smoking when *b+1* years old or later, or 4) formerly smoked but stopped smoking when *b-2* years old or earlier. Smokers included: 1) current smokers who started smoking when *b-1* years old or earlier, and 2) former smokers who started smoking when *b-1* years old or early but stopped smoking when *b+1* years old or later.

*Potential confounders*

The age of participants at baseline was derived by the UK Biobank based on their date of birth and date of attending the initial assessment. We used Townsend deprivation index and household income as measures of socio-economic position at baseline. Townsend deprivation index is a score measuring material deprivation in the area where participants were living, calculated by UK Biobank using participants' postcode. Participants (except those who living in sheltered accommodation or a care home) were asked to report their average total household income before tax, with five categories ranging from less than £18,000 to greater than £100,000. Sex of participants was acquired from the registry and updated by the participants.

Supplementary Table 5 summarised the UK Biobank fields we used to derive our smoking phenotypes, outcomes, and confounders.

### SUPPLEMENTARY FIGURES

**Supplementary Figure 1. Participant flow diagram**

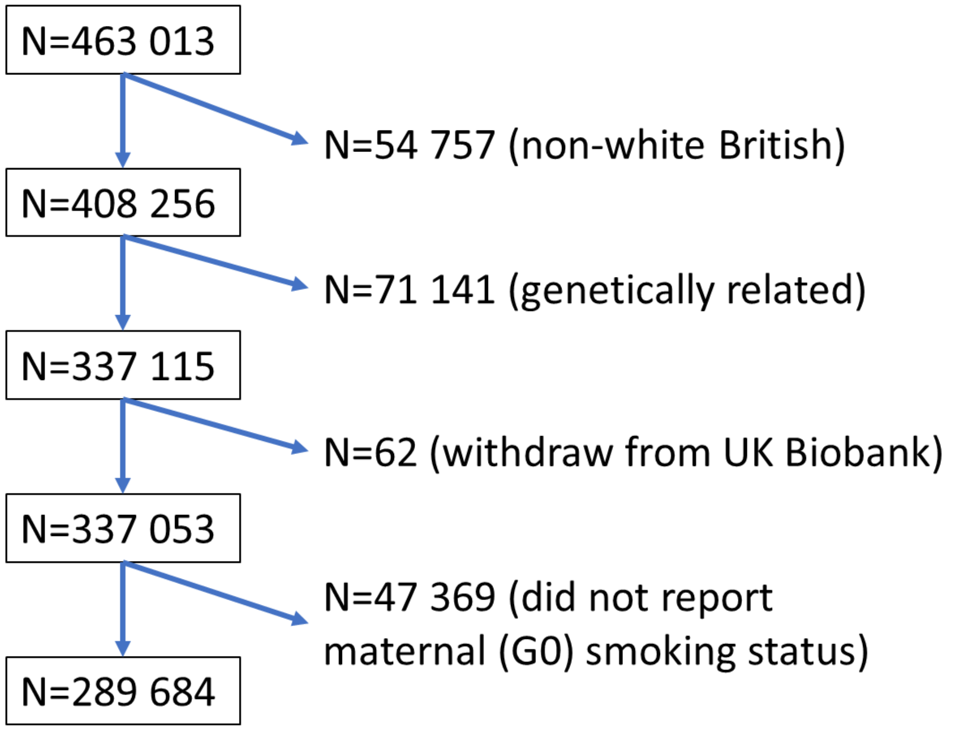

Note: The 463,013 participants are of European descent, have genetic sex same as reported sex, either XX or XY in sex chromosome, and no outliers in heterozygosity and missing rates (1).

**Supplementary Figure 2. Directed acyclic graphs illustrating potential collider bias**

(A)

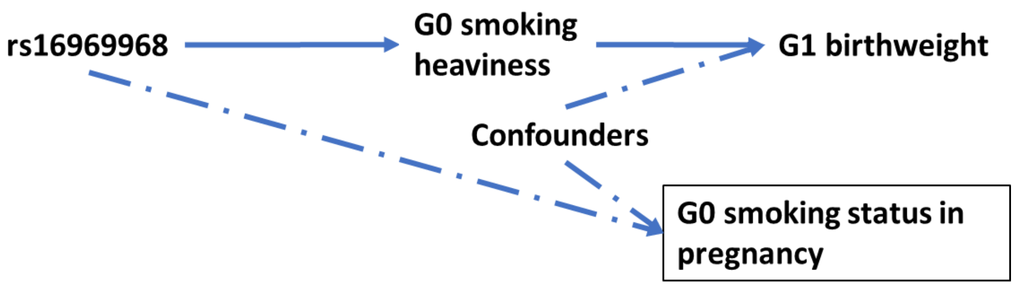

(B)

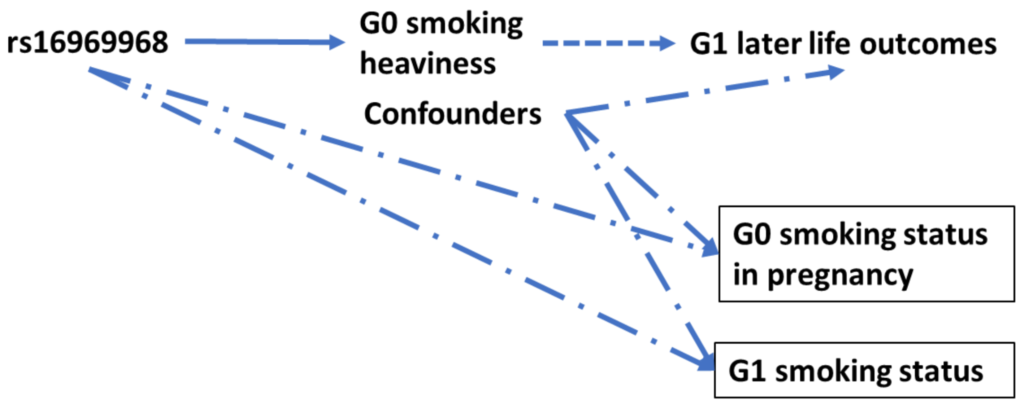

(C)

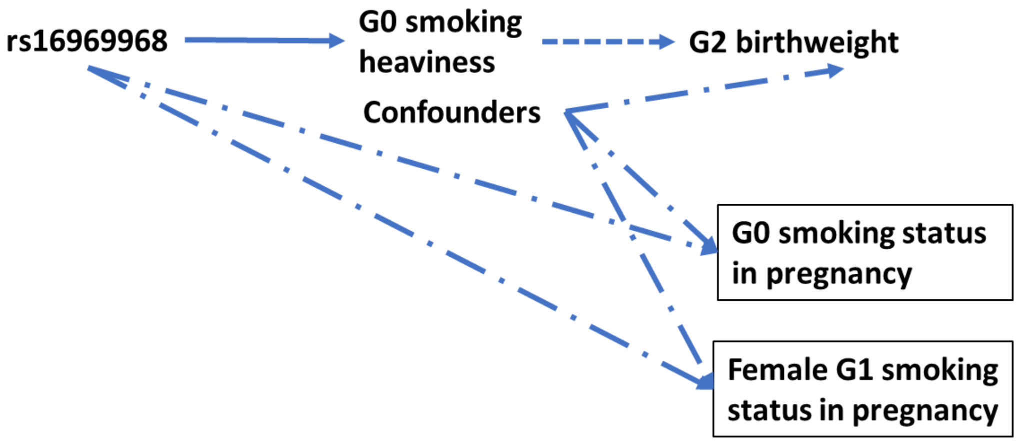

Generation (G)0: UK Biobank participants' mother; G1: UK Biobank participants themselves; G2: First offspring of UK Biobank participants. Boxes around a phenotype denote this phenotype is being conditioned upon. Solid arrow denotes known association, dashed arrow denotes the hypothesis we are testing, and the dashed-dotted arrow denotes potential alternative pathways via a collider.

(A) We stratified maternal (G0) smoking status in pregnancy (shown in rectangle as it is conditioned on) in our proof of principle analysis. The smoking heaviness variant (i.e. rs16969968) was known to influence smoking status in pregnancy (2), and it was verified in our study. Our stratification may induce collider bias between rs16969968 and G1 birthweight, if there are common causes between G0 smoking status and G1 birthweight (i.e. there is an unblocked path between them). We would have an unblocked path (shown as — · ▸ ) between rs16969968 and G1 birthweight that is not via G0 smoking heaviness.

(B) Besides the potential unblocked path described in (A), we further stratified G1 participants on their own smoking status, which rs16969968 is also associated with in our study, when we estimated the effect on G1 later life outcomes (shown in - - - ▸ ). This may induce more collider bias between rs16969968 and G1 later life outcomes due to another unblocked path via conditioning on G1 smoking status (shown as — · ▸ ).

(C) Besides the potential unblocked path described in (A), we further stratified G1 women on their smoking status in pregnancy, when we explored the effect on G2 birthweight (shown in - - - ▸ ). Due to the same reason in (A), we may have another unblocked path between rs16969968 and G2 birthweight (shown as — · ▸ ).

Other sources of bias regarding conditioning on exposure or outcomes (e.g. due to missingness) have been discussed in reference (3).

**Supplementary Figure 3. The associations of rs16969968 with UK Biobank participants height and age at menarche in sensitivity analyses**

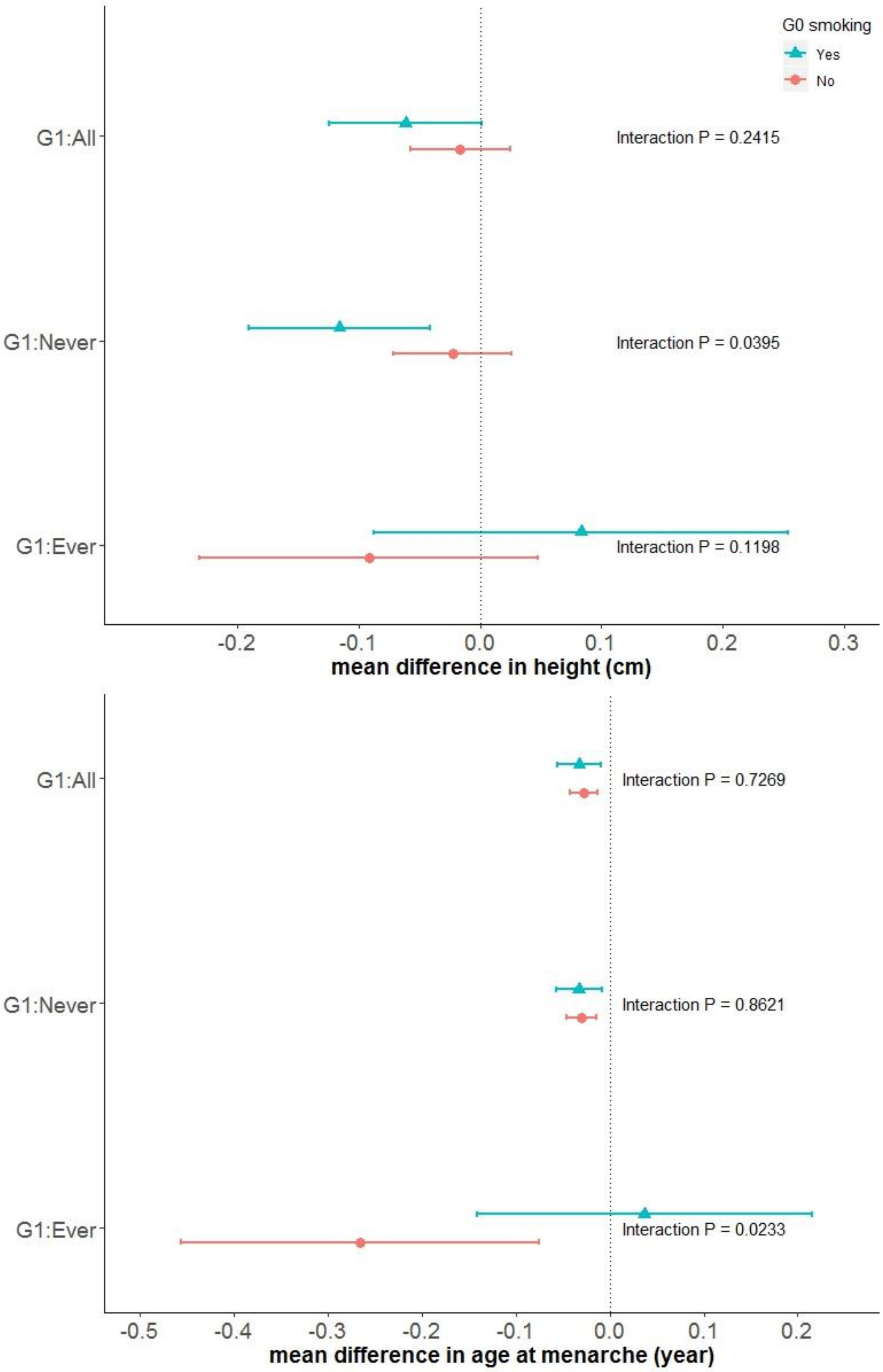

Generation (G)0: UK Biobank participants' mother; G1: UK Biobank participants themselves. Estimates are the mean difference of G1 outcome per each smoking-heaviness increasing allele of rs16969968. G1 were grouped according to whether they were ever smokers before achieving their adulthood height or their age at menarche. G1 who started smoking at the same age of achieving their adulthood height or at menarche were removed from analyses due to uncertainty.

**Supplementary Figure 4. Illustration of edge cases when deriving participants smoking in pregnancy**

**A) Participants started smoking in the same year as the first live birth (N=538)**

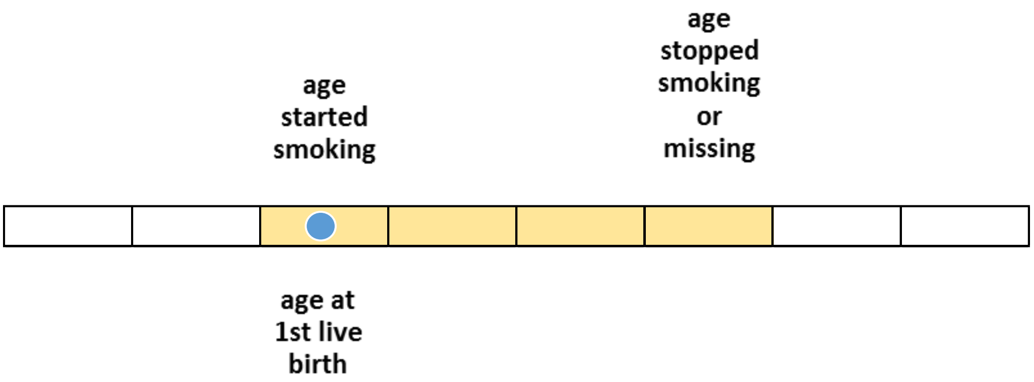

**B) Participants stopped smoking in the same year as the first live birth (N=1129)**

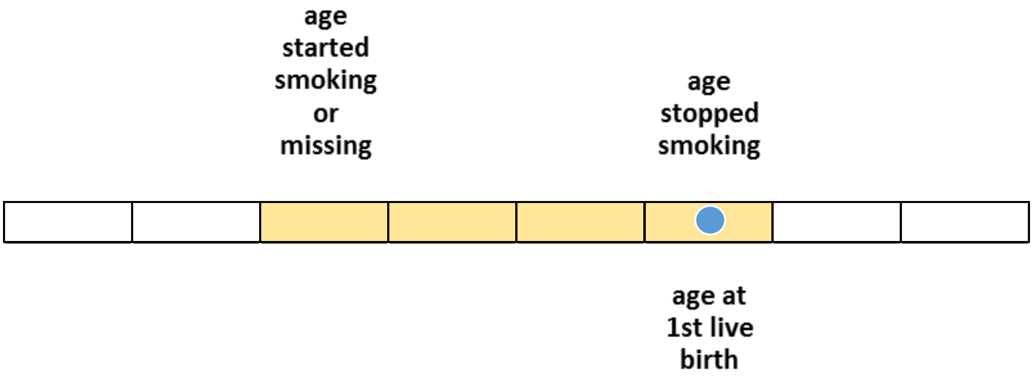

**C) Participants stopped smoking in the year before the first live birth (N=1017)**

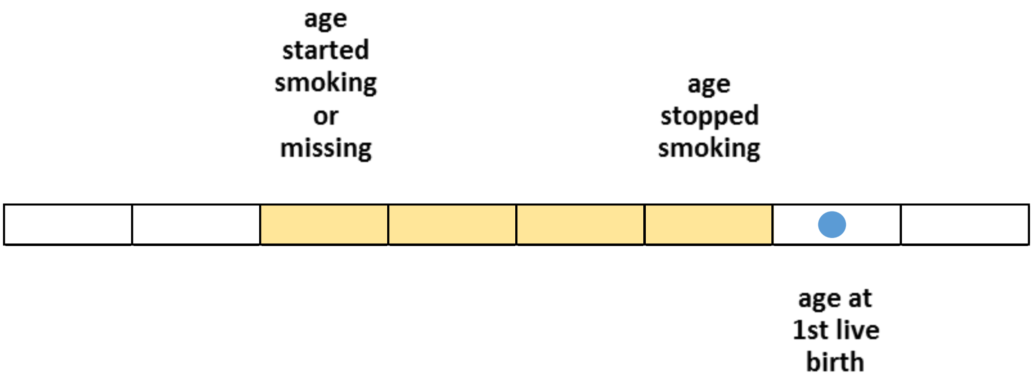

This figure illustrates our derivation of female participants smoking status in their pregnancy resulting in their first live birth, described in the Supplementary Methods section above.

### SUPPLEMENTARY TABLES

**Supplementary Table 1. Characteristics of the UK Biobank participants by sex**

| Variable | | Total N | Sex [1] | |
|---|---|---|---|---|
| | | | Men | Women |
| ***Smoking*** | | | | |
| Maternal smoking in pregnancy | *Yes* | 88 447 | 41 791 (31.5) | 46 656 (29.7) |
| | *No* | 201 237 | 90 889 (68.5) | 110 348 (70.3) |
| Smoking status of participants | *Current* | 27 420 | 14 924 (11.3) | 12 496 (8.0) |
| | *Former* | 99 932 | 51 209 (38.7) | 48 723 (31.1) |
| | *Never* | 161 398 | 66 108 (50.0) | 95 290 (60.9) |
| Participants smoking in pregnancy | *Yes* | 19 061 | Not applicable | 19 061 (19.1) |
| | *No* | 80 552 | | 80 552 (80.9) |
| ***Outcome*** | | | | |
| Participants' birthweight (kg) | | 171 784 | 3.45 ± 0.68 | 3.24 ± 0.63 |
| Standing height (cm) | | 289 050 | 176.04 ± 6.75 | 162.77 ± 6.22 |
| Body mass index ($kg/m^2$) | | 288 775 | 27.79 ± 4.21 | 26.98 ± 5.12 |
| Forced vital capacity (L) | | 218 378 | 4.48 ± 0.87 | 3.21 ± 0.63 |
| Forced expiratory volume in 1-second (L) | | 218 378 | 3.37 ± 0.73 | 2.44 ± 0.52 |
| Asthma | *Case* | 30 751 | 13 008 (9.8) | 17 743 (11.3) |
| | *Control*[2] | 258 639 | 119 531 (90.2) | 139 108 (88.7) |
| Age at menarche (year) | | 152 991 | Not applicable | 12.95 ± 1.60 |
| Systolic blood pressure (mmHg) | | 289 430 | 141.15 ± 17.44 | 135.33 ± 19.13 |
| Diastolic blood pressure (mmHg) | | 289 431 | 84.14 ± 10.00 | 80.64 ± 9.95 |
| Year of Education | | 287 198 | 15.44 ± 5.06 | 14.63 ± 5.05 |
| Fluid intelligence score | | 148 462 | 6.17 ± 2.18 | 5.95 ± 2.03 |
| Depression/anxiety | *Case* | 102 637 | 36 333 (27.4) | 66 304 (42.3) |
| | *Control* | 186 743 | 96 197 (72.6) | 90 546 (57.7) |
| Happiness | *Extremely happy* | 5417 | 2727 (6.2) | 2690 (5.2) |
| | *Very happy* | 37 872 | 17 670 (40.1) | 20 202 (39.2) |
| | *Moderately happy* | 48 223 | 21 618 (49.0) | 26 605 (51.6) |
| | *Moderately unhappy* | 3358 | 1677 (3.8) | 1681 (3.3) |
| | *Very unhappy* | 566 | 283 (0.6) | 283 (0.5) |
| | *Extremely unhappy* | 188 | 108 (0.2) | 80 (0.2) |
| Birthweight of female participants' first child (kg) | | 126 122 | Not applicable | 3.19 ± 0.55 |
| ***Potential confounder*** | | | | |
| Age (year) | | 289 684 | 56.97 ± 8.12 | 56.48 ± 7.95 |
| Age at first live birth (years) | | 107 277 | Not applicable | 25.48 ± 4.55 |
| Deprivation index | | 289 334 | -1.59 ± 2.96 | -1.64 ± 2.86 |
| Household income | *less than £18,000* | 52 806 | 22 437 (18.7) | 30 369 (23.1) |
| | *£18,000 to £30,999* | 63 807 | 29 163 (24.3) | 34 644 (26.4) |
| | *£31,000 to £51,999* | 66 999 | 32 882 (27.4) | 34 117 (26.0) |
| | *£52,000 to £100,000* | 53 588 | 27 876 (23.2) | 25 712 (19.6) |
| | *greater than £100,000* | 14 235 | 7659 (6.4) | 6576 (5.0) |

[1] Mean ± standard deviation for continuous variables and N (column %) for categorical variables.

[2] Controls were participants who did not indicate having asthma diagnosed by a doctor.

**Supplementary Table 2. Associations of rs16969968 with potential confounders by strata of smoking status in participants (G1) and their mothers (G0), adjusted for the first ten principal components**

| Confounder | By G1 smoking status | Overall | By G0 smoking status in pregnancy | |
|---|---|---|---|---|
| | | | Yes | No |
| Age (years)[1] | All participants | -0.018 (-0.062, 0.026) | -0.080 (-0.156, -0.004) | 0.013 (-0.040, 0.067) |
| | Current smokers | -0.073 (-0.218, 0.072) | -0.092 (-0.325, 0.141) | -0.061 (-0.245, 0.124) |
| | Former smokers | -0.077 (-0.149, -0.006) | -0.044 (-0.170, 0.083) | -0.087 (-0.174, -0.001) |
| | Ever smokers | -0.089 (-0.154, -0.024) | -0.062 (-0.175, 0.051) | -0.097 (-0.176, -0.017) |
| | Never smokers | 0.054 (-0.005, 0.113) | -0.074 (-0.177, 0.028) | 0.113 (0.041, 0.185) |
| | Female participants | 0.013 (-0.046, 0.072) | -0.059 (-0.163, 0.045) | 0.049 (-0.022, 0.121) |
| | Female smokers in pregnancy | -0.159 (-0.315, -0.003) | 0.037 (-0.215, 0.290) | -0.291 (-0.487, -0.095) |
| | Female non-smokers in pregnancy | 0.055 (-0.026, 0.136) | -0.077 (-0.220, 0.067) | 0.121 (0.023, 0.220) |
| Age at first live birth (years)[1] | Female participants | 0.019 (-0.022, 0.060) | 0.050 (-0.024, 0.125) | 0.015 (-0.034, 0.063) |
| | Female smokers in pregnancy | 0.015 (-0.072, 0.101) | 0.010 (-0.128, 0.148) | 0.017 (-0.093, 0.127) |
| | Female non-smokers in pregnancy | 0.024 (-0.026, 0.074) | 0.068 (-0.026, 0.163) | 0.017 (-0.041, 0.076) |
| Townsend deprivation index[1] | All participants | 0.010 (-0.006, 0.026) | 0.002 (-0.029, 0.032) | 0.011 (-0.007, 0.030) |
| | Current smokers | 0.018 (-0.042, 0.079) | 0.019 (-0.084, 0.123) | 0.018 (-0.056, 0.093) |
| | Former smokers | 0.008 (-0.019, 0.035) | -0.014 (-0.065, 0.038) | 0.015 (-0.017, 0.047) |
| | Ever smokers | 0.016 (-0.010, 0.041) | -0.002 (-0.050, 0.045) | 0.022 (-0.008, 0.051) |
| | Never smokers | 0.012 (-0.008, 0.032) | 0.014 (-0.024, 0.052) | 0.009 (-0.014, 0.032) |
| | Female participants | -0.002 (-0.023, 0.019) | -0.018 (-0.059, 0.023) | 0.002 (-0.022, 0.027) |
| | Female smokers in pregnancy | -0.026 (-0.092, 0.040) | -0.098 (-0.208, 0.011) | 0.021 (-0.062, 0.103) |
| | Female non-smokers in pregnancy | -0.002 (-0.029, 0.026) | -0.022 (-0.076, 0.031) | 0.002 (-0.029, 0.034) |
| Years of education[1] | All participants | 0.036 (0.008, 0.064) | 0.045 (-0.006, 0.097) | 0.036 (0.003, 0.069) |
| | Current smokers | 0.030 (-0.064, 0.124) | 0.092 (-0.065, 0.249) | -0.006 (-0.123, 0.111) |
| | Former smokers | 0.043 (-0.005, 0.091) | 0.022 (-0.068, 0.112) | 0.055 (-0.003, 0.112) |
| | Ever smokers | 0.038 (-0.006, 0.081) | 0.038 (-0.040, 0.117) | 0.040 (-0.012, 0.091) |

| | | | | |
|---|---|---|---|---|
| | Never smokers | 0.026 (-0.010, 0.062) | 0.036 (-0.032, 0.103) | 0.026 (-0.017, 0.069) |
| | Female participants | 0.050 (0.012, 0.088) | 0.064 (-0.006, 0.133) | 0.049 (0.005, 0.094) |
| | Female smokers in pregnancy | 0.035 (-0.074, 0.144) | 0.155 (-0.022, 0.332) | -0.038 (-0.176, 0.100) |
| | Female non-smokers in pregnancy | 0.046 (-0.006, 0.098) | 0.095 (-0.002, 0.191) | 0.033 (-0.029, 0.095) |
| Household income[2] | All participants | 0.996 (0.986, 1.007) | 0.992 (0.974, 1.011) | 0.998 (0.986, 1.011) |
| | Current smokers | 0.979 (0.947, 1.013) | 0.980 (0.926, 1.037) | 0.978 (0.937, 1.021) |
| | Former smokers | 0.993 (0.975, 1.010) | 0.995 (0.963, 1.028) | 0.992 (0.971, 1.013) |
| | Ever smokers | 0.988 (0.972, 1.004) | 0.990 (0.962, 1.018) | 0.987 (0.969, 1.006) |
| | Never smokers | 0.999 (0.985, 1.013) | 0.988 (0.963, 1.013) | 1.003 (0.987, 1.020) |
| | Female participants | 0.994 (0.980, 1.009) | 0.991 (0.965, 1.018) | 0.996 (0.979, 1.014) |
| | Female smokers in pregnancy | 1.000 (0.959, 1.043) | 1.012 (0.946, 1.082) | 0.991 (0.940, 1.046) |
| | Female non-smokers in pregnancy | 1.006 (0.986, 1.027) | 1.004 (0.967, 1.042) | 1.008 (0.984, 1.033) |
| Sex (compare to female)[3] | All participants | 0.995 (0.984, 1.006) | 0.987 (0.967, 1.007) | 0.998 (0.985, 1.011) |
| | Current smokers | 0.994 (0.959, 1.030) | 0.992 (0.935, 1.052) | 0.995 (0.951, 1.041) |
| | Former smokers | 1.004 (0.985, 1.023) | 1.005 (0.972, 1.040) | 1.003 (0.981, 1.026) |
| | Ever smokers | 1.002 (0.986, 1.019) | 1.003 (0.973, 1.033) | 1.002 (0.982, 1.023) |
| | Never smokers | 0.993 (0.978, 1.008) | 0.977 (0.951, 1.004) | 0.999 (0.981, 1.017) |

[1] Results were from linear regression. Estimates are mean difference of confounder per each smoking-heaviness increasing allele of rs16969968.

[2] Results were from ordinal logistic regression for household income (less than £18 000 = 1, £18 000 to £30 999 = 2, £31 000 to £51 999 = 3, £52 000 to £100 000 = 4, greater than £100 000 = 5). Estimates are the change in odds of being in a higher level of household income per each smoking-heaviness increasing allele of rs16969968.

[3] Results from logistic regression for sex. Estimates are the change in odds of being male rather than female per each smoking-heaviness increasing allele of rs16969968.

**Supplementary Table 3. Differences in the associations of rs16969968 with 12 outcomes in participants (G1) across maternal (G0) smoking status in pregnancy**

| G1 outcome (associations shown in Figure 3) | By G1 smoking status[1] | Interaction P-value[2] |
|---|---|---|
| Height | All participants | 0.2415 |
| | Never smokers | 0.0291 |
| | Ever smokers | 0.5491 |
| Body mass index | All participants | 0.8710 |
| | Never smokers | 0.4258 |
| | Former smokers | 0.2756 |
| | Current smokers | 0.7849 |
| Forced expiratory volume in 1-second | All participants | 0.4603 |
| | Never smokers | 0.6462 |
| | Former smokers | 0.4970 |
| | Current smokers | 0.7575 |
| Forced vital capacity | All participants | 0.6788 |
| | Never smokers | 0.3199 |
| | Former smokers | 0.9120 |
| | Current smokers | 0.1879 |
| Asthma | All participants | 0.9756 |
| | Never smokers | 0.7751 |
| | Ever smokers | 0.8507 |
| Systolic blood pressure | All participants | 0.3985 |
| | Never smokers | 0.2725 |
| | Former smokers | 0.7075 |
| | Current smokers | 0.5221 |
| Diastolic blood pressure | All participants | 0.5861 |
| | Never smokers | 0.9896 |
| | Former smokers | 0.1164 |
| | Current smokers | 0.2265 |
| Age at menarche | All participants | 0.7269 |
| | Never smokers | 0.8938 |
| | Ever smokers | 0.8250 |
| Years of education | All participants | 0.9796 |
| | Never smokers | 0.8282 |
| | Ever smokers | 0.8791 |
| Fluid intelligence score | All participants | 0.9067 |
| | Never smokers | 0.2581 |
| | Former smokers | 0.1856 |
| | Current smokers | 0.8321 |
| Depression/anxiety | All participants | 0.8368 |
| | Never smokers | 0.8355 |
| | Former smokers | 0.9112 |
| | Current smokers | 0.6190 |
| Happiness | All participants | 0.0668 |
| | Never smokers | 0.1529 |
| | Ever smokers | 0.2689 |

[1] We combined current and former smokers into ever smokers for some outcomes given smoking cessation may not influence them rapidly.

[2] Interaction P-value was obtained using Cochran's Q statistic for the heterogeneity in the association of rs16969968 with each outcome between participants whose mothers did versus did not smoke.

**Supplementary Table 4. Observational associations of participants mothers' smoking in pregnancy with participants' smoking status and all outcomes**

| Dependent variables | Model 1 | | Model 2 | |
|---|---|---|---|---|
| ***Smoking*** | *OR (95% CI)* | *P-value* | | |
| Smoking status of participants (ever vs never) | 1.073 (1.056, 1.090) | $3.7\times10^{-18}$ | Not applicable | |
| Participants smoking in pregnancy (smoker vs non-smoker) | 1.494 (1.446, 1.544) | $1.3\times10^{-126}$ | Not applicable | |
| ***Outcome*** | *Mean difference (95% CI)* | *P-value* | *Mean difference (95% CI)* | *P-value* |
| Participants' birthweight (kg)[1] | -0.104 (-0.111, -0.097) | $6.6\times10^{-198}$ | -0.101 (-0.107, -0.094) | $1.1\times10^{-182}$ |
| Standing height (cm)[1] | -0.509 (-0.582, -0.437) | $8.2\times10^{-43}$ | -0.273 (-0.345, -0.200) | $1.6\times10^{-13}$ |
| Body mass index (kg/m²)[1] | 0.850 (0.813, 0.888) | ~0 | 0.732 (0.695, 0.770) | $1.9\times10^{-319}$ |
| Forced vital capacity (L)[1] | -0.037 (-0.045, -0.028) | $4.7\times10^{-17}$ | -0.010 (-0.018, -0.001) | $2.3\times10^{-2}$ |
| Forced expiratory volume in 1-second (L)[1] | -0.038 (-0.044, -0.031) | $4.0\times10^{-29}$ | -0.015 (-0.022, -0.008) | $7.4\times10^{-6}$ |
| | *OR (95% CI)* | *P-value* | *OR (95% CI)* | *P-value* |
| Asthma (case vs control)[1] | 1.074 (1.047, 1.101) | $4.2\times10^{-8}$ | 1.075 (1.048, 1.103) | $3.5\times10^{-8}$ |
| | *Mean difference (95% CI)* | *P-value* | *Mean difference (95% CI)* | *P-value* |
| Age at menarche (year)[1] | -0.056 (-0.074, -0.039) | $3.5\times10^{-10}$ | -0.070 (-0.087, -0.052) | $1.3\times10^{-14}$ |
| Systolic blood pressure (mmHg)[1] | 0.517 (0.377, 0.656) | $3.7\times10^{-13}$ | 0.436 (0.296, 0.577) | $1.2\times10^{-9}$ |
| Diastolic blood pressure (mmHg)[1] | 0.413 (0.333, 0.493) | $5.4\times10^{-24}$ | 0.393 (0.312, 0.474) | $1.9\times10^{-21}$ |
| Year of Education[1] | -0.796 (-0.836, -0.757) | ~0 | Not applicable | |
| Fluid intelligence score[1] | -0.236 (-0.259, -0.212) | $1.1\times10^{-87}$ | -0.132 (-0.154, -0.110) | $5.2\times10^{-31}$ |
| | *OR (95% CI)* | *P-value* | *OR (95% CI)* | *P-value* |
| Depression/anxiety (case vs control)[1] | 1.191 (1.172, 1.211) | $4.9\times10^{-97}$ | 1.160 (1.141, 1.179) | $5.3\times10^{-68}$ |
| Happiness (extremely happy to extremely unhappy)[1] | 0.940 (0.915, 0.965) | $4.8\times10^{-6}$ | 0.958 (0.932, 0.984) | $1.7\times10^{-3}$ |
| | *Mean difference (95% CI)* | *P-value* | *Mean difference (95% CI)* | *P-value* |
| Birthweight of female participants' first child(kg)[2] | -0.003 (-0.009, 0.004) | 0.44 | 0.013 (0.005, 0.020) | $6.2\times10^{-4}$ |

[1] Model 1 adjusted for participants' age; Model 2 adjusted for participants' age, smoking status, education and Townsend deprivation index.

[2] Model 1 adjusted for participants' age; Model 2 adjusted for participants' age, smoking status in pregnancy, education and Townsend deprivation index.

**Supplementary Table 5. UK Biobank data field of variables used in this study**

| Variable | UK Biobank data field |
|---|---|
| ***Smoking phenotypes*** | |
| Maternal smoking around birth | 1787 |
| Smoking status | 20116 |
| Age started smoking in current smokers | 3436 |
| Age started smoking in former smokers | 2867 |
| Age stopped smoking | 2897 |
| ***Outcomes*** | |
| Birth weight | 20022 |
| Standing height | 50 |
| Body mass index (BMI) | 21001 |
| Forced expiratory volume in 1-second (FEV1), Best measure | 20150 |
| Forced vital capacity (FVC), Best measure | 20151 |
| Blood clot, DVT, bronchitis, emphysema, asthma, rhinitis, eczema, allergy diagnosed by doctor | 6152 |
| Systolic blood pressure, automated reading | 4080 |
| Systolic blood pressure, manual reading | 93 |
| Diastolic blood pressure, automated reading | 4079 |
| Diastolic blood pressure, manual reading | 94 |
| Age when periods started (menarche) | 2714 |
| Qualifications | 6138 |
| Fluid intelligence score | 20016, 20191 |
| Seen doctor (GP) for nerves, anxiety, tension or depression | 2090 |
| Seen a psychiatrist for nerves, anxiety, tension or depression | 2100 |
| Diagnoses - main ICD10 | 41202 |
| Diagnoses - secondary ICD10 | 41204 |
| Happiness | 4526 |
| Birth weight of first child | 2744 |
| ***Potential confounder*** | |
| Age at recruitment | 21022 |
| Age at first live birth | 2754 |
| Sex | 31 |
| Townsend deprivation index at recruitment | 189 |
| Average total household income before tax | 738 |

Further information on these phenotypes can be found by searching the UK Biobank data showcase: http://biobank.ctsu.ox.ac.uk/showcase/search.cgi.

**References**

1.Mitchell R, Hemani G, Dudding T, Paternoster L. UK Biobank Genetic Data: MRC-IEU Quality Control, Version 1. 06 Nov 2017. https://data.bris.ac.uk/data/dataset/3074krb6t2frj29yh2b03x3wxj.

2.Freathy RM, Ring SM, Shields B*, et al.* A common genetic variant in the 15q24 nicotinic acetylcholine receptor gene cluster (CHRNA5-CHRNA3-CHRNB4) is associated with a reduced ability of women to quit smoking in pregnancy. *Hum Mol Genet* 2009;18:2922-7. doi: 10.1093/hmg/ddp216

3.Hughes RA, Davies NM, Davey Smith G, Tilling K. Selection bias in instrumental variable analyses. *BioRxiv* 2018. doi: 10.1101/192237
